# Endogenous cell competition kills pathogenic *de novo* mutant cells but declines under pH stress

**DOI:** 10.64898/2026.04.27.721036

**Authors:** Yuki Akieda, Hikari Ukawa, Natsuki Watanabe, Tohru Ishitani

## Abstract

Although endogenous cell competition safeguards embryonic development by eliminating unfit cells, the intrinsic origins of unfitness remain unclear. Here, zebrafish whole-genome sequencing and imaging reveal that endogenous cell competition removes cells with spontaneous *de novo* mutations in key developmental signalling pathways and in human disease-associated genes, such as those associated with Alzheimer’s disease, epilepsy, autism, and premature aging. These mutations arise frequently during normal development but are efficiently eliminated under physiological conditions, thereby preventing the clonal expansion of harmful cells. However, environmental pH stress disrupts this surveillance by dysregulating Ca^2+^ signals, allowing mutant cells to persist and resulting in tissue mispatterning and morphological defects. Our findings establish endogenous cell competition as an intercellular communication-dependent genome quality-control mechanism that removes pathogenic mutant cells before clonal expansion. This link between developmental robustness, environmental stress, and mosaic genetic disorders highlights how intrinsic mutations and extrinsic stressors together shape disease susceptibility across the lifespan.

## Introduction

Embryonic development requires precise coordination of signalling networks, metabolic programs, and morphogenetic processes to form correctly patterned tissues. Remarkably, embryos achieve this process with high reproducibility despite substantial perturbations. This characteristic, known as developmental robustness, was first conceptualized by Conrad Waddington^1,2^. Various mechanisms actively detect and correct intrinsic errors that inevitably arise during development, such as stochastic fluctuations in gene expression, metabolic imbalances, or signal transduction errors. One key candidate mechanism for error-correction is cell competition, a quality control process in which unfit cells are selectively eliminated based on their differences in cellular fitness^3^. Since its discovery, cell competition has been primarily studied as a cell-cell communication-dependent mechanism that eliminates artificially introduced abnormal cells, including those with mutations in ribosome and oncogenic genes, in *Drosophila* imaginal discs and mammalian cell cultures^4–11^. Additionally, growing evidence from mouse and zebrafish embryos suggests that “endogenous cell competition” is active during normal development, eliminating cells with unfit morphogen signalling (e.g., Wnt or Shh pathways)^12,13^, impaired mitochondrial activity^14^, reduced pluripotency due to low YAP-TEAD activity^15^, or low Myc activity^16^. However, the intrinsic origins of such cellular unfitness, including the fundamental triggers of endogenous cell competition, remain unclear.

Here, we show that during normal embryogenesis, pathogenic *de novo* mutant cells — including mutations in key signalling pathways and in human disease–associated genes such as those linked to Alzheimer’s disease, epilepsy, autism spectrum disorder (ASD), and premature aging—are spontaneously generated and trigger endogenous cell competition. Under physiological conditions, endogenous cell competition acts as a genome surveillance system to eliminate these pathogenic cells. However, this safeguard is disrupted by environmental pH stress, allowing mutant cells to persist. Although these results establish endogenous cell competition as a critical safeguard against intrinsic developmental errors, this safeguard can decline under extrinsic stress, revealing a hidden vulnerability within otherwise robust development.

## Results

### Cell competition eliminates *de novo* mutant cells during development

Although cell competition can eliminate cells with various unfitness during physiological development^3^, the intrinsic origins of such cellular unfitness remain unclear. Identifying these origins is essential for improving the understanding of cell competition and development. Given the rapidity of the cell cycle and DNA replication during embryogenesis, we hypothesized that genomic errors, including mutations and chromosomal aberrations, arise frequently and may trigger cell competition–mediated elimination. To test this hypothesis, we first examined DNA double-strand breaks (DSBs) in embryos. DSBs cause mutations, insertions/deletions, and chromosomal loss during repair^17–19^. Even in normally developing zebrafish embryos, cells positive for the DSB marker γH2AX (γH2AX^+^) were detected (Fig. 1A, left panel), suggesting that cells with DSBs naturally arise during physiological development. To examine whether cells with DSBs and consequent genomic errors are eliminated through apoptosis-mediated cell competition, we inhibited this process by overexpressing selenophosphate synthetase1 (*sephs1*), a negative regulator of cell competition, and the anti-apoptotic factor B-cell lymphoma-2 (*bcl-2*)^12^. Overexpression of *sephs1* and *bcl-2* mRNA led to the persistence of γH2AX^+^ cells (Fig. 1A and B), suggesting that cells harbouring DSBs, which frequently give rise to mutations or chromosomal abnormalities during repair, are ultimately eliminated through competition. DSBs can lead to severe chromosomal aberrations in rapidly dividing early human and mouse embryos^17,18^. Therefore, we investigated whether inhibition of cell competition, which allows cells harbouring DSBs to persist, leads to chromosomal abnormalities in embryos. We found that inhibition of cell competition led to the accumulation of cells with micronuclei and nuclear buds, which are indicators of chromosomal abnormalities (Fig. 1C and D). Additionally, nuclei containing micronuclei were merged with γH2AX (Fig. 1E). These results suggest that although cells with DSBs, which can cause severe genomic abnormalities, arise during normal development, endogenous cell competition acts to eliminate these cells and preserves the genomic integrity of embryos, thereby contributing to developmental robustness.

**Fig. 1.**
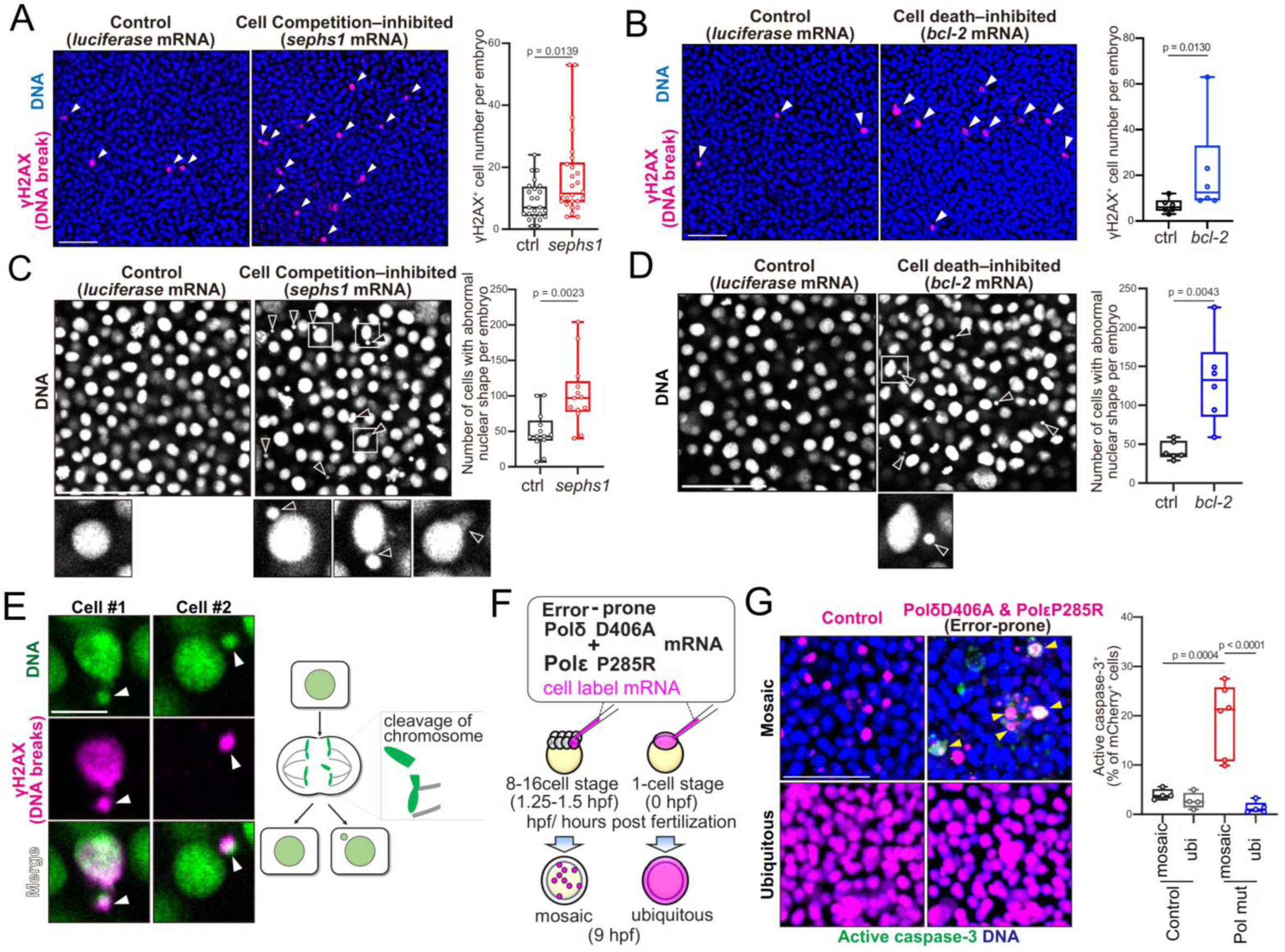
Cell competition eliminates *de novo* mutant cells during development. **(A,B)** In cell competition– and cell death–inhibited embryos, cells with DNA double-strand breaks are increased. Confocal images showing whole-mount immunostaining of γH2AX and DNA in *sephs1* (**A**) and *bcl-2* mRNA–injected embryos (**B**). Scale bars, 50 μm. Arrowheads indicate γH2AX^+^ cells. Box plots show the number of γH2AX^+^ cell per embryo on the dorsal side; each dot represents one embryo. (**C, D)** In cell competition– and cell death–inhibited embryos, cells with abnormal nuclear morphology are increased. Confocal images show whole-mount Hoechst33342 staining in *sephs1* (**C**) and *bcl-2* mRNA–injected embryos (**D**), with magnified views. Scale bars, 50 μm. Arrowheads indicate nuclear abnormalities, such as micronuclei. Box plots show the number of cells with abnormal nuclear morphology per embryo on the dorsal side; each dot represents one embryo. **(E)** Abnormal nuclei merged with γH2AX (DNA break). Confocal images show immunostaining of γH2AX and DNA in *sephs1* mRNA–injected embryos. Scale bars, 10 μm. (**F)** Experimental design for introducing error-prone DNA polymerase mutants into zebrafish embryos. Polδ D406A and Polε P285R mRNAs were co-injected with a fluorescent nuclear maker at the 8-16-cell stage (1.25-1.5 hpf) to generate mosaic mutant cells, or at the one-cell sage (0 hpf) to generate ubiquitous mutants. Embryos were fixed at 9 hpf for analysis (**G)** Cells expressing error-prone polymerase mutants undergo apoptosis in a cell competition–dependent manner. Confocal images showing immunostaining of active caspase-3 in mosaic or ubiquitous embryos expressing H2B-mCherry with/without Polδ and Polε mutants. Scale bars, 50 μm. Box plots show the percentage of mCherry^+^ cells that are active caspase-3^+^, calculated by dividing the number of double-positive cells by the total number of mCherry^+^ cells; each dot represents one embryo. In total, approximately 400-1,200 label^+^ cells were quantified per condition across embryos.

To confirm that cell competition eliminates cells with *de novo* mutations, we introduced the error-prone polymerase δ (Polδ) D406A and polymerase ε (Polε) P285R mutants into zebrafish embryos in a mosaic or ubiquitous manner (Fig. 1F). These Polδ/Polε mutants lack proofreading activity and accumulate mutations at a high rate. In transgenic mice, such mutations lead to a high mutational burden, ultimately resulting in cancer^20,21^. Mosaically introduced Polδ D406A and Polε P285R mutants induced apoptosis, whereas ubiquitous introduction did not (Fig. 1G), suggesting that neighbouring normal cells were necessary for eliminating artificially introduced genome-mutated cells. These results suggest that early embryonic tissues can eliminate cells with *de novo* mutations through a cell competition–dependent mechanism.

### Cell competition eliminates pathogenic *de novo* mutant cells

To identify *de novo* gene mutations that are eliminated by endogenous cell competition during embryogenesis, we performed whole-genome sequencing (WGS) of pooled, apoptosis-inhibited zebrafish larvae, in which *de novo* mutant cells were expected to accumulate (Fig. 2A). This approach preferentially detects mutations present in clonally expanded cell populations that have persisted to the larval stage. The results revealed 760 mutations across various genes in apoptosis-inhibited larvae but not in control larvae. Among these, 104 were classified as high-impact mutations, including frameshift and splice-site mutations, whereas 566 were categorized as moderate-impact mutations, involving single or multiple amino acid substitutions (Fig. 2B). In contrast, we did not detect clear differences in structural variants, including gene fusions, or copy number variations (CNVs) between control larvae and apoptosis-inhibited larvae under the experimental conditions used. Gene Ontology analysis of these mutations demonstrated enrichment of signal transduction–related gene sets (Fig. 2C). Notably, some of these mutated genes have also been implicated in driving cell competition. Consistent with our previous report that cells with abnormal Wnt signalling are eliminated through cell competition^12^, we detected single-nucleotide substitution mutations, frameshift mutations, and splice acceptor site mutations caused by insertions/deletions in genes encoding Wnt signalling molecules (Fig. 2D). Specifically, frameshift mutations in the Wnt receptor frizzled/FZD (*fzd3b*)^22^, amino acid substitution mutations in the Wnt co-receptor family LRP1 (*lrp1aa* and *lrp1bb*)^23,24^, and splice acceptor site mutations in the Wnt agonist R-spondin (*rspo4*)^25,26^ were detected. These genes regulate canonical Wnt signalling^22–25^, and knockout or mutations in these genes result in severe developmental abnormalities^23,27–30^. These findings suggest that during normal physiological development, *de novo* mutations occur in genes involved in Wnt signalling, and such mutant cells are eliminated via apoptosis mediated by endogenous cell competition.

**Fig. 2.**
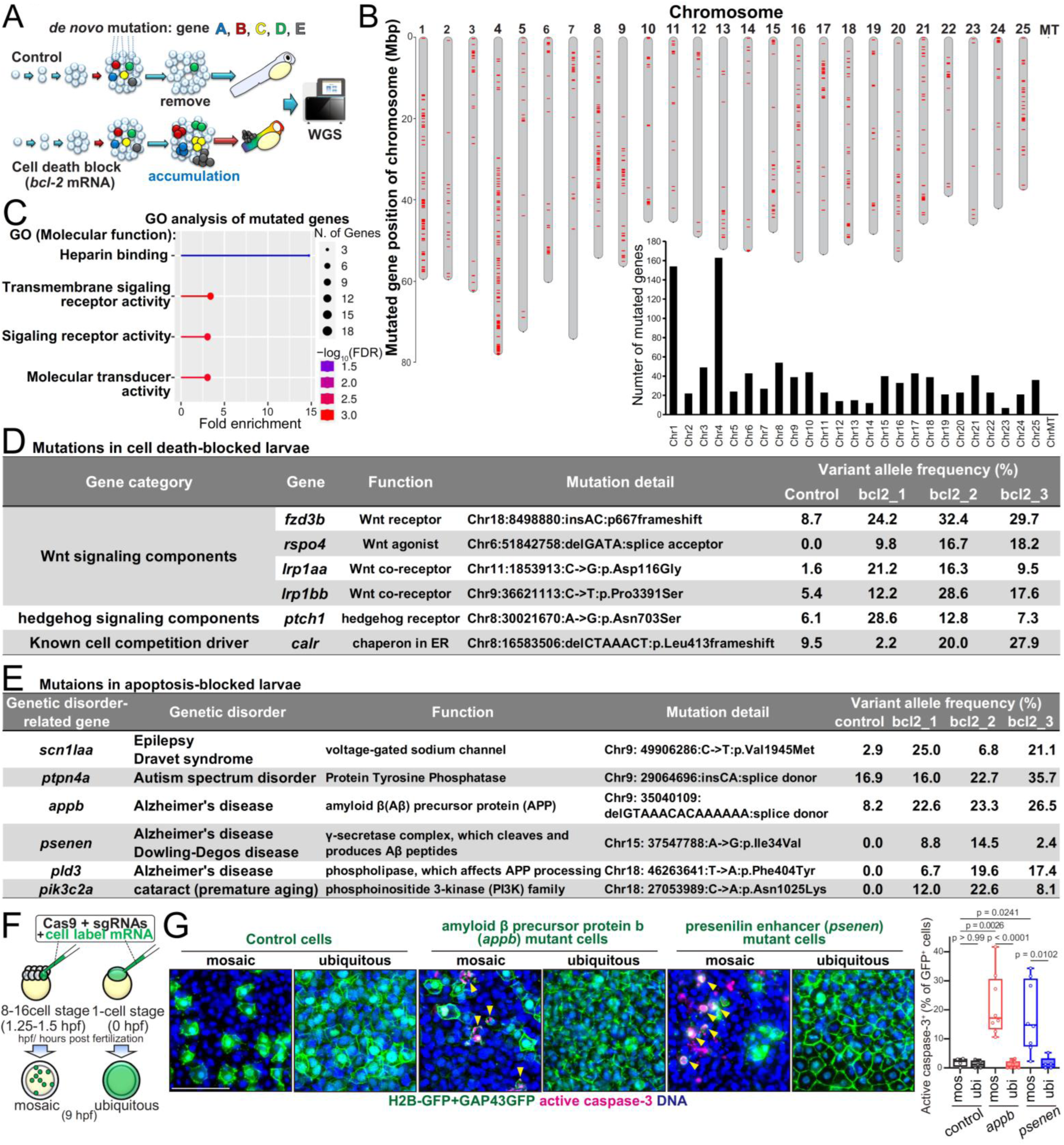
Endogenous cell competition eliminates pathogenic *de novo* mutant cells during embryogenesis. **(A)** Experimental overview. *de novo* mutations arise during normal development and are removed by cell competition (control), whereas blocking cell death (*bcl-2* mRNA–injected) causes their accumulation, enabling detection by whole-genome sequencing (WGS). The illustration of next-generation sequencing is from TogoTV (© 2016 DBCLS TogoTV, CC-BY-4.0). (**B**) Chromosomal distribution of mutated genes identified in apoptosis-inhibited larvae. Red bars indicate mutated loci. The graph (bottom) shows the number of mutated genes per chromosome. (**C)** Gene Ontology (molecular function) analysis of mutated genes detected in apoptosis-inhibited larvae. (**D)** Representative mutations in genes previously linked to cell competition. The table lists mutated genes involved in Wnt/hedgehog signalling or the known competition-related factor *calr*, with variant allele frequencies in each sample. VAF (variant allele frequency) indicates the proportion of sequencing reads carrying a given mutation relative to total reads at that genomic site, reflecting the mosaic frequency of mutant cells in the sample. (**E**) Mutations in genes associated with human mosaic genetic disorders. The table lists representative genes (eg, *scn1a*, *ptpn4*, *appb*, *psenen*, *pld3*, *pik3c2a*) detected in apoptosis-inhibited larvae, with VAF. (**F**) Experimental strategy for introducing mutant cells were generated using Cas9 and sgRNAs, injected at the 8-16-cell stage (1.25-1.5 hpf) for mosaic induction or at the one-cell stage (0 hpf) for ubiquitous induction. Embryos were fixed at 9 hpf for analysis. (**G**) Amyloid β precursor protein b (*appb*) mutant and presenilin enhancer (*psenen*) mutant cells undergo apoptosis in a cell competition–dependent manner. Images show whole-mount immunostaining of active caspase-3 in mosaic or ubiquitous embryos introduced label^+^ control, *appb* or *psenen* mutant cells. Box plots show the percentage of label^+^ cells that are active caspase-3^+^, calculated by dividing the number of double-positive cells by the total number of label^+^ cells; each dot represents one embryo. In total, approximately 300-700 label^+^ cells were quantified per condition across embryos.

In addition to morphogen Wnt signalling, amino acid substitution mutations in patched1 (*ptch1*), a receptor for morphogen sonic hedgehog (Shh) signalling, were detected (Fig. 2D). Artificially introduced cells with abnormal Shh signalling in mouse skin epithelial tissues and zebrafish larvae were also reported to drive cell competition^13,31^. In addition to the signalling pathways, a frameshift mutation in calreticulin (*calr*), an endoplasmic reticulum chaperone molecule that drives cell competition in the *Drosophila* eye imaginal disc^32^, was detected (Fig. 2D).

Furthermore, recent next-generation sequencing studies demonstrated that *de novo* mutations occur in early human embryos^33,34^. Genetic mosaicism arising from *de novo* mutations during embryogenesis has been implicated in various disorders, such as epilepsy, ASD, and Alzheimer’s disease (*35*–*37*). Unexpectedly, WGS analysis of zebrafish larvae with inhibited apoptosis revealed mutations in genes associated with various diseases linked to mosaicism. Specifically, we detected mutations in the SCN1A (sodium channel protein type 1 subunit alpha), PTCH1, PTPN4 (protein tyrosine phosphatase non-receptor type-4), APP (amyloid β precursor protein), PSENEN (presenilin enhancer γ-secretase subunit), PLD3 (phospholipase D3), and PIK3C2A (phosphatidylinositol-4-phosphate 3-kinase catalytic subunit type 2 alpha) genes, corresponding to *scn1laa*, *ptch1*, *ptpn4a*, *appb*, *psenen*, *pld3*, and *pik3c2a* in zebrafish, respectively (Fig. 2E). SCN1A is the causative gene of epilepsy and Dravet syndrome, a severe form of epilepsy in infants^35,36^. Mosaic mutations in this gene have been detected in the brains of epilepsy patients, and SCN1A-mutant zebrafish exhibit epileptic seizures^37^. PTCH1 *de novo* mosaic mutations cause hypothalamic hamartoma with gelastic epilepsy^35^. PTPN4 mosaic mutations were identified in patients with ASD^38^. APP^39^, PSENEN^40^, and PLD3^41^ are causative genes of Alzheimer’s disease and are involved in APP processing. Notably, cells artificially expressing amyloid β in *Drosophila* can be eliminated through cell competition^42^. In addition, PSENEN mutations are responsible for Dowling–Degos disease, an autosomal-dominant disorder of skin pigmentation. Knockdown of *psenen* in zebrafish larvae induces phenotypes that closely mimic the human manifestation of this disease^43^. PIK3C2A mutations cause premature aging symptoms and the early onset of cataracts in humans, mice, and zebrafish^44^. Therefore, these results suggest that although pathogenic *de novo* mutant cells carrying disease-associated mutations naturally arise, such cells are usually eliminated through endogenous cell competition during embryogenesis.

To confirm whether cell competition eliminates *de novo* mutant cells associated with these mosaic genetic disorders, we introduced mutant cells into zebrafish embryos using the CRISPR/Cas9 system in a mosaic or ubiquitous manner (Fig. 2F). Mosaically introduced amyloid β precursor protein b (*appb*) mutant cells or presenilin enhancer (*psenen*) mutant cells underwent apoptosis, whereas ubiquitous introduction did not (Fig. 2G), suggesting that cells harbouring mutations in Alzheimer’s disease-associated genes are eliminated by endogenous cell competition. These results also suggest that cell competition prevents mosaic genetic disorders by eliminating pathogenic *de novo* mutant cells.

### Environmental pH stress disrupts cell competition

Our findings suggest that endogenous cell competition acts as a critical safeguard against “intrinsic developmental errors (*de novo* mutations)” to ensure developmental robustness. However, some “extrinsic perturbations (environmental stresses)” can affect tissue development and homeostasis regardless of robustness. For example, ocean acidification causes abnormal development in marine organisms^45–47^, and maternal diabetes during pregnancy causes body fluid acidification and diverse fetal malformations^48–50^, similar to the phenotypes observed in zebrafish larvae, which inhibit cell competition^12^. Therefore, we hypothesized that external perturbation, particularly pH stress, interferes with cell competition (Fig. 3A). Consistent with our previous observations, artificially introduced cells with abnormal Wnt signalling, which express constitutively active β-catenin (βcatCA; Wnt-High) or GSK-3β (Wnt-Low), were apoptotically eliminated from zebrafish embryos and produced no gross morphological abnormalities in a normal water environment with pH 7 (Fig. 3B and C and Fig. S1A). However, acidic (pH 6; one unit below normal) and alkaline (pH 8; one unit above normal) environments inhibited apoptotic elimination and caused malformations (Fig. 3B and C and Fig. S1A–C). Milder acidic environments (pH 6.5; 0.5 unit below normal) and alkaline environments (pH 7.7; 0.7 unit above normal) also blocked apoptotic elimination of cells with abnormal Wnt signalling (Fig. S1D), suggesting that cell competition is ineffective at pH levels below 6.5 or above 7.7. We also confirmed that, in addition to hydrochloric acid, acidic environments prepared with lactic acid and acetic acid, which are abundant in the tumour microenvironment, suppressed apoptosis of cells with abnormal Wnt signalling (Fig. S1E). Furthermore, both acidic and alkaline pH stresses suppressed the apoptotic elimination of cells expressing Polδ and Polε mutants, and *appb* mutant (Fig. 3B, D, and E), suggesting that pH stress suppresses the elimination of *de novo* mutant cells. Remarkably, pH stress alone did not cause malformations in the majority of individuals (Fig. S1C). This observation is consistent with a recent study which reported that environmental acidification does not cause overt lethality or widespread apoptosis^51^. Thus, our results suggest that environmental pH stress disrupts the elimination of abnormal cell via cell competition but does not significantly affect the main system that drives the developmental program.

**Fig. 3.**
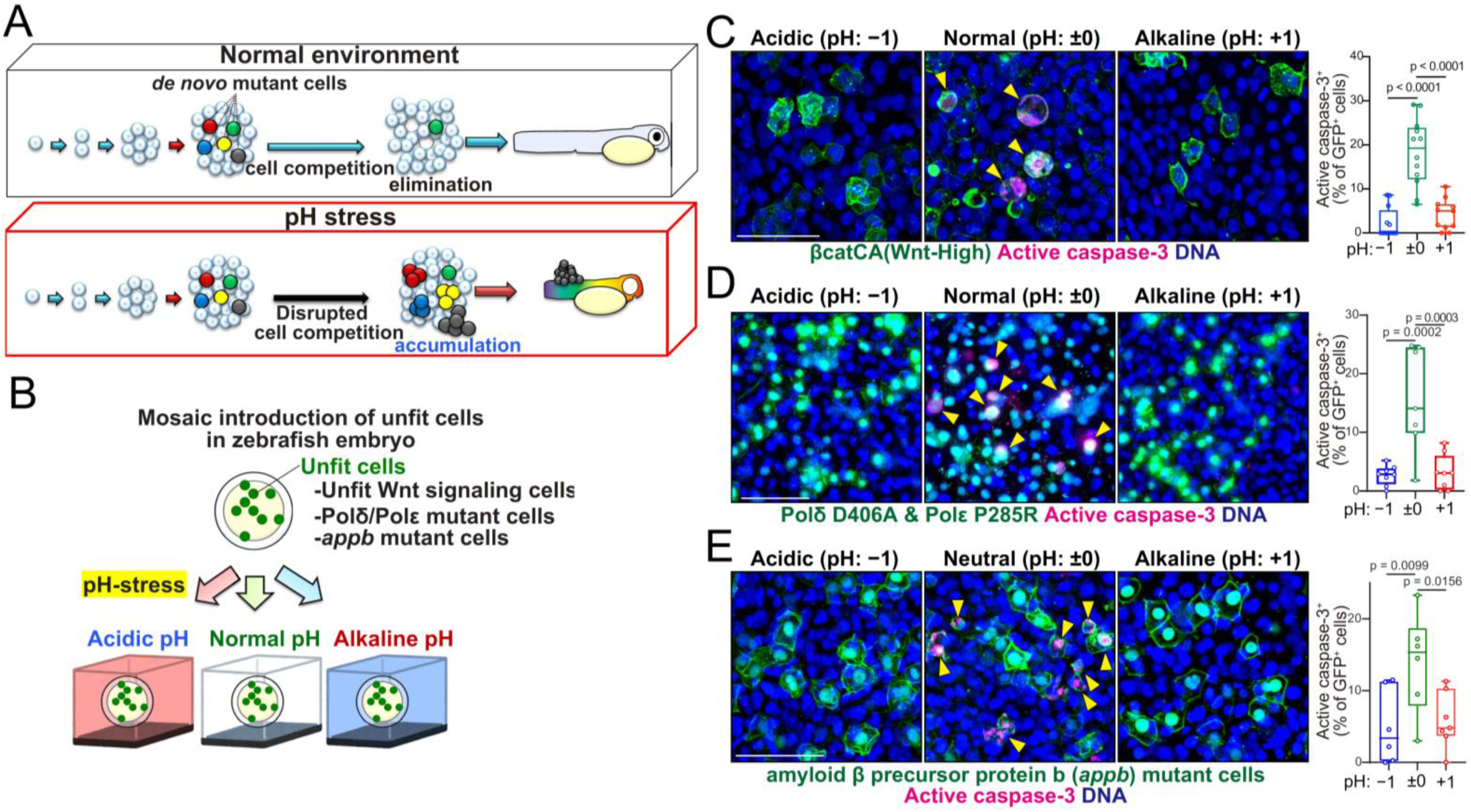
Environmental pH stress disrupts cell competition. **(A)** Schematic illustration of environmental pH stress disrupting cell competition. (**B**) Experimental design. Fluorescently labelled unfit cells (unfit Wnt signalling cells, error-prone Polδ/ε mutant-expressing cells, *appb* mutant cells) were mosaically introduced into zebrafish embryos and then subjected to pH stress. (**C**) Artificially introduced βcatCA-expressing (Wnt-High) cells undergo apoptosis under normal pH but not under acidic and alkaline pH. Confocal images show whole-mount immunostaining of active caspase-3 and DNA. Scale bars, 50 μm. Arrowheads indicate βcatCA-expressing active caspase-3^+^ cells. Box plots show the percentage of βcatCA-expressing cells (Wnt-High) that are active caspase-3^+^, calculated by dividing the number of double-positive cells by the total number of βcatCA-expressing cells; each dot represents one embryo. In total, approximately 300-400 βcatCA-expressing cells were quantified per condition across embryos. (**D**) Artificially introduced error-prone Pol mutant-expressing cells undergo apoptosis in a normal pH environment but not in acidic and alkaline pH environments. Images show whole-mount immunostaining of active caspase-3 and DNA. Scale bars, 50 μm. Arrowheads indicate Pol mutant-expressing active caspase-3^+^ cells. Box plots show the percentage of Pol mutant-expressing cells that are active caspase-3^+^, calculated by dividing the number of double-positive cells by the total number of Pol mutant-expressing cells; each dot represents one embryo. In total, approximately 900-1,000 Pol mutant-expressing cells were quantified per condition across embryos. (**E**) Amyloid β precursor protein (*appb*) mutant cells undergo apoptosis in a normal pH environment but not in acidic and alkaline pH environments. Images show whole-mount immunostaining of active caspase-3 and DNA. Scale bar, 50 μm. Arrowheads indicate *appb* mutant active caspase-3^+^ cells. Box plots show the percentage of *appb* mutant cells that are active caspase-3^+^, calculated by dividing the number of double-positive cells by the total number of *appb* mutant cells; each dot represents one embryo. In total, approximately 300-800 *appb* mutant cells were quantified per condition across embryos.

### pH stress disrupts cell competition–mediated developmental robustness

To confirm whether pH stress also suppresses endogenous cell competition, we measured the numbers of cells with DSBs, apoptotic cells, and cells with abnormal nuclei. In a normal water environment at pH 7, both γH2AX^+^ cells with DSBs and active caspase-3^+^ apoptotic cells were detected (Fig. 4A and B). These γH2AX^+^ cells are likely to be eliminated by endogenous cell competition. As expected, both acidic and alkaline pH stresses caused a decrease in apoptotic cells in normal embryos (Fig. 4B) and an increase in γH2AX^+^ and micronuclei^+^ cells (genome-error cells) (Fig. 4A and C). Our previous study showed that inhibition of cell competition leads to the accumulation of cells with ectopic activation and abnormal inactivation in the Wnt signalling gradient, thereby causing the accumulation of mispatterned cells and improper embryonic morphogenesis^12^. Hence, we investigated whether disruption of cell competition via pH stress also caused these abnormalities. In embryos exposed to pH stress, cells with ectopically high or abnormally low Wnt signalling accumulated (Fig. 4D). Furthermore, the number of mispatterned cells aberrantly expressing the Wnt signalling target posterior gene *tbx6* increased (Fig. 4E). For instance, *tbx6*-expressing anterior cells and *tbx6*-lacking posterior cells were detected (Fig. 4E). Consistent with these results, pH stress during embryogenesis impaired normal muscle segment formation (Fig. 4F), which is precisely controlled via *tbx6* expression^52^. These results suggest that pH stress disrupts endogenous cell competition and interferes with correct tissue patterning. Thus, pH stress-mediated disruption of endogenous cell competition causes only minor abnormalities, such as the accumulation of a few abnormal cells. However, as development progresses, embryos exposed to pH stress exhibited an increase in the number of abnormal individuals, in which one eye was small, one pectoral fin was small, or a tumour-like cell mass had formed over time (Fig. 4G and H). Similar defects were observed in *sephs1* mRNA–injected cell competition–inhibited embryos (Fig. 4G and H). These abnormalities were likely caused by gradual proliferation and differentiation of the few surviving abnormal cells. Thus, pH stress disrupts endogenous cell competition, leading to pathological cell accumulation that may negatively impact future body quality.

**Fig. 4.**
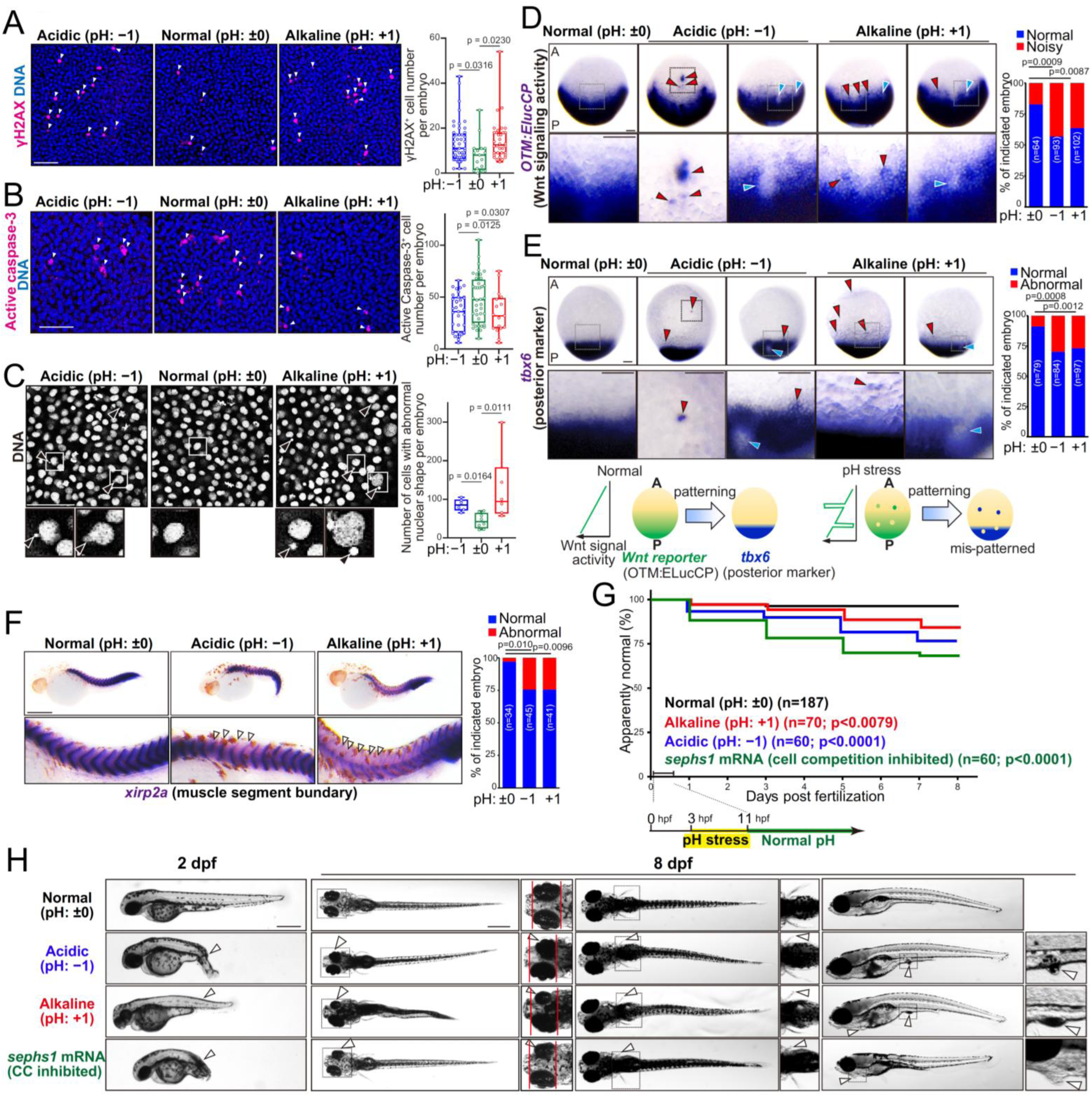
pH stress disrupts cell competition–mediated developmental robustness. (**A–C**) pH stress increases endogenous cells with *de novo* mutations and reduces apoptosis. Confocal images showing whole-mount immunostaining of anti-γH2AX (**A**), anti-active caspase-3 (**B**), and Hoechst33342/DNA (**C**) in embryos under normal, acidic, or alkaline pH. Scale bar, 50 μm. Arrowheads indicate γH2AX^+^ cells, active caspase-3^+^ cells, and nuclear abnormalities (e.g., micronuclei). Box plots show the number of γH2AX^+^ cell, active caspase-3^+^, and cells with nuclear abnormalities per embryo on the dorsal side; each dot represents one embryo. (**D, E**) pH stress promotes abnormal Wnt signalling cells and tissue mispatterning. Whole-mount *in situ* hybridization showing ELuc-CP (Wnt reporter) in Tg(Optimal TCF Motif/OTM:ELuc-CP) embryos (**D**, dorsal view) and *tbx6* (posterior marker) in embryos (**E**, lateral view, dorsal to the right) under the same pH conditions. A, anterior; P, posterior. Scale bars, 200 μm. Percentages of embryos with normal or abnormal morphology are indicated. (**F**) Whole-mount *in situ* hybridization of *xirp2a* (muscle segment boundary) in larvae at 28 hpf. Scale bars, 500 μm. Percentages of larvae with normal or abnormal morphology are shown. (**G**) Graph shows the rate of increase in post-fertilization malformations in zebrafish treated at normal, acidic, or alkaline pH during embryonic development (3–11 hpf) or injected with *sephs1* mRNA. (**H**) Representative images show malformed phenotypes from (**G**) at 2 and 8 dpf. Arrowheads indicate abnormal morphology. Scale bars, 500 μm.

### pH stress causes chaotic Ca^2+^ influx to disrupt cell competition

Finally, we investigated the mechanism by which pH stress disrupts cell competition. At acidic pH, measurements using zebrafish embryos expressing a pH sensor (mCherry-SEpHluorin^53^) showed that the pH of the extracellular interstitial fluid shifted substantially, whereas the intracellular pH changed slightly compared to the extracellular pH (Fig. S2A). This result suggests that environmental pH stress affects the extracellular space. Additionally, both acidic and alkaline pH environments induce chaotic Ca^2+^ influx from extracellular to intracellular space through certain ion channels including mechano-sensitive channels^54–56^. Our previous studies showed that during Wnt/Shh-unfitness-driven cell competition, Wnt/Shh-unfit cells alter membrane cadherin levels to generate an imbalance in cadherin levels between unfit and neighbouring normal cells. This cadherin imbalance leads to local changes in mechanical tension, which then activates the PIEZO family of mechano-sensitive channels and consequent Ca^2+^ influx specifically in adjacent normal cells, triggering recognition and elimination of unfit cells (Fig. 5A, left panel)^12,57^. Similarly, mosaically generated *app* mutant cells decreased membrane E-cadherin levels and generated a cadherin imbalance between mutant and neighbouring normal cells (Fig. S2B), indicating that *app* mutant cells are eliminated through the Ca^2+^ influx-mediated mechanism similar to Wnt/Shh-unfitness-driven cell competition (*58*). Based on our previous findings and these results, pH stress, which can cause dysregulated Ca^2+^ influx, may interfere with the proper function of cell competition (Fig. 5A, right panel). We first confirmed that both acidic and alkaline pH stress induced chaotic Ca^2+^ influx in normal zebrafish embryos by using the Ca^2+^ probe GCaMP7 (Fig. 5B and C and movie S1). To determine whether dysregulated Ca^2+^ influx inhibits cell competition, we forcibly induced Ca^2+^ influx from the extracellular to intracellular space by treating embryos with ionomycin, a calcium ionophore (Fig. S2C and movie S2). Ionomycin treatment suppressed apoptotic elimination of cells with abnormally high Wnt signalling (βcatCA) and *appb* mutant cells (Fig. 5D and E). Thus, pH stress disrupts cell competition through chaotic Ca^2+^ influx.

**Fig. 5.**
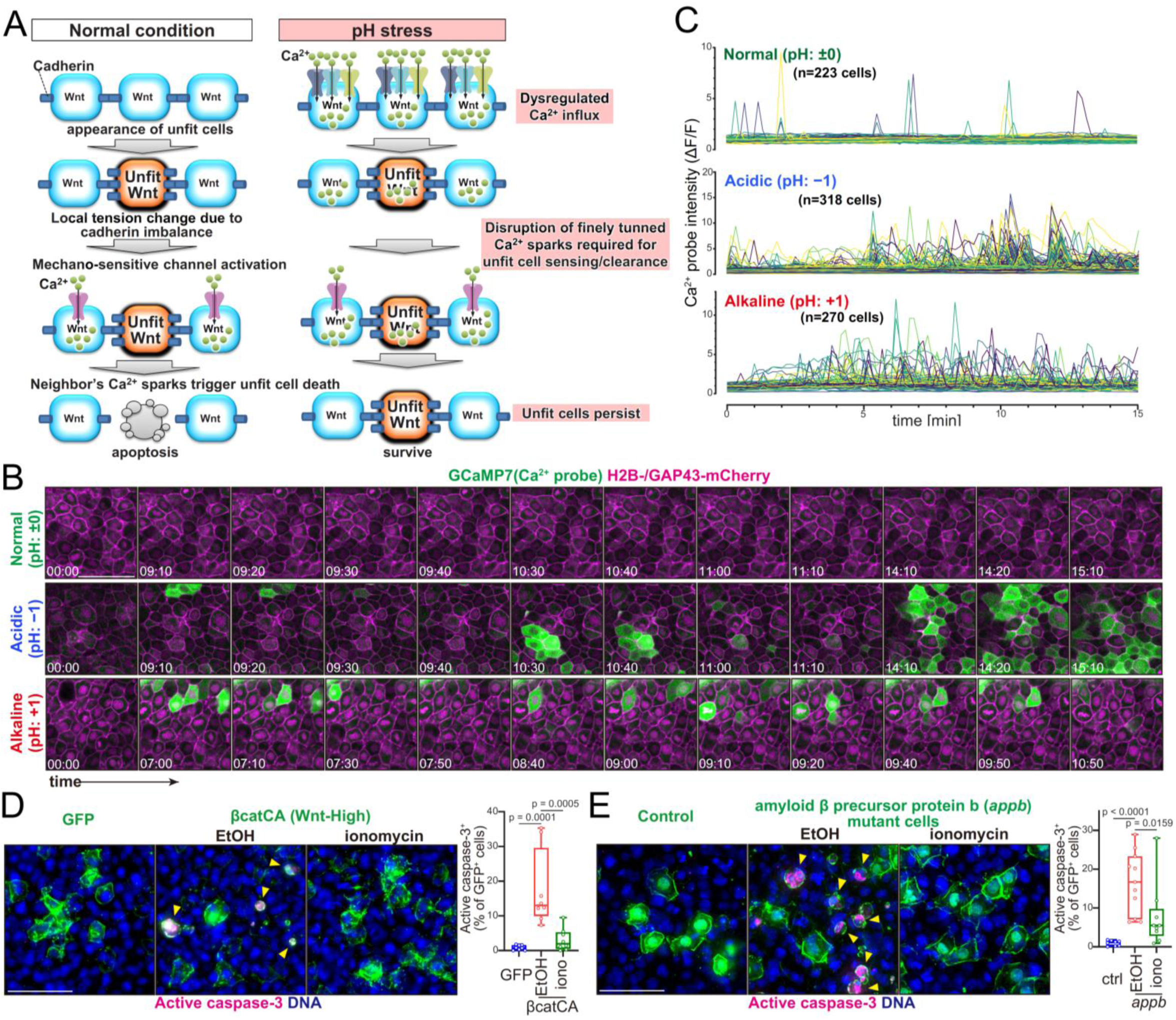
pH stress disrupts cell competition through chaotic Ca^2+^ influx. (**A**) Schematic illustration of the mechanisms underlying Wnt-driven cell competition. (**B**) pH stress induces dysregulated Ca^2+^ influx. Time-lapse confocal fluorescence images of deep cells in embryos co-expressing the calcium probe GCaMP7 and nuclear/membrane-localized mCherry (H2B-/GAP43-mCherry) under normal, acidic, or alkaline pH. Scale bar, 50 μm. (**C**) Graphs show traces of Ca^2+^ probe fluorescence intensity in deep cells and EVL cells over 15 min. Each line represents an individual cell. (**D, E**) Forced Ca^2+^ influx inhibits cell competition. Confocal images show whole-mount immunostaining of active caspase-3 and DNA in embryos mosaically expressing βcatCA (**D**) or amyloid β precursor protein b (*appb*) mutant cells (**E**), treated with ethanol or ionomycin. Scale bar, 50 μm. Arrowheads indicate active caspase-3⁺ mutant cells. Box plots show the percentage of labelled cells that are active caspase-3^+^, calculated by dividing the number of double-positive cells by the total number of labelled cells; each dot represents one embryo. In total, approximately 2,000 βcatCA-expressing cells, or approximately 2,800-6,200 *appb* mutant cells were quantified per condition across embryos.

## Discussion

In this study, we demonstrate that endogenous cell competition serves as a physiological surveillance mechanism that eliminates *de novo* mutant cells arising spontaneously during normal embryogenesis. Endogenous cell competition eliminates cells with mutations not only in genes regulating key developmental signalling pathways but also in genes implicated in human disease such as Alzheimer’s disease, epilepsy, ASD and premature aging. This clearance process contributes to developmental robustness and protects the embryo from latent disease risk. Importantly, we found that environmental pH stress disrupts this surveillance by inducing chaotic Ca^2+^ influx, thereby preventing the recognition and elimination of mutant cells. As a result, mutant cells persist and accumulate, potentially leading to mosaic genetic disorders later in life. This finding provides a mechanistic link between developmental robustness and the embryonic origins of mosaic genetic disorders, including neurodevelopmental and neurodegenerative diseases.

Cell competition has been widely studied as a fitness-sensing mechanism that removes unfit cells in various contexts, including differences in protein synthesis^11,58^, Myc expression^16,59,60^, YAP-TEAD activity^15^, and morphogen Wnt/Shh signalling^12,13,61^. However, a fundamental question in cell competition research—the intrinsic origins of cellular unfitness that trigger endogenous cell competition—has remained unclear. We found that spontaneously arising *de novo* mutations are a substantial source of such unfitness, particularly within Wnt/Shh signalling genes and endoplasmic reticulum chaperones associated with protein synthesis. Notably, a recent study using adult mouse epidermis reported a conceptually related tissue-level quality control mechanism, which selectively eliminates stem cells harbouring DSBs through differentiation induction, accompanied by compensatory clonal expansion of neighbouring intact stem cells, thereby safeguarding genomic integrity^62^. Moreover, experimentally induced aneuploid cells are selectively eliminated through cell competition in mouse embryos^63,64^, human pluripotent stem cells^65^, and *Drosophila* imaginal disc^66^. Extending beyond these artificial systems and DNA damage–based models, our study demonstrates that cells carrying pathogenic *de novo* mutations are likewise recognized and eliminated by apoptosis-mediated endogenous cell competition during embryogenesis. Together, these findings reframe cell competition not merely as a modulator of relative cellular fitness, but as a broader genome surveillance system that safeguards genomic integrity during development.

Physiological pH is tightly regulated, with extracellular pH (e.g., blood pH) normally maintained at 7.4 and intracellular pH at 7.2. However, in pathological contexts such as tumour microenvironments, ischemic tissues, or inflammatory sites, extracellular pH can fall below 6.5^67–70^. In these contexts, metabolic alterations substantially contribute to acidification. Similarly, studies on ocean acidification have shown that in marine organisms such as sea urchins and clownfish, even modest reductions in seawater pH from the normal 8.0 to 7.6–7.8 can disrupt embryonic development and organ formation^45–47^. In our study, even relatively mild changes in pH (pH 6.5; 0.5 unit below normal or pH 7.7; 0.7 unit above normal) tend to suppress cell competition. Taken together, these findings suggest that cell competition is vulnerable to pH stress within physiologically and pathologically relevant ranges, providing a mechanism by which acidosis interferes with developmental robustness and tissue homeostasis. More broadly, a recent study reported that metabolic stresses can also modulate cell competition. For example, extracellular L-proline deprivation suppresses unfit cell elimination via integrated stress response^71^. Because metabolic states are coupled to pH homeostasis, these observations suggest that nutritional and pH stresses may converge to weaken cell competition. These insights support the view that cell competition may be broadly vulnerable to diverse environmental perturbations.

Maternal diabetes during pregnancy increases the risk of fetal malformations. The phenotypes of malformations are diverse and include caudal regression, neural tube defects, heterotaxia (abnormal arrangement across the left-right axis of the body), widely variable fetal size, and cardiovascular abnormalities^49,50,72^. Several mechanisms have been proposed as the cause of malformations, including epigenetic changes^73^, oxidative stress^50^, and abnormal Nodal signalling due to high glucose levels^72^. This study suggests a previously unidentified mechanism linking diabetes and fetal malformations: diabetic acidosis may induce the accumulation of cells with *de novo* mutations by abrogating cell competition. Our results revealed that reduced cell competition caused the accumulation of cells with mutations in genes encoding Wnt or Shh signalling regulators, which control caudal development, neural tube formation, and body size. Wnt signalling is required for tail bud formation, and *Wnt3a* knockout mice show caudal regression^74^. Shh signalling also influences neural tube patterning and body size; *ptch1* knockout mice show neural tube defects, and transgenic mice highly expressing *ptch1* exhibit approximately 50% smaller body sizes^75–77^. Therefore, under pH stress, such as acidosis, cells with *de novo* mutations in the Wnt and Shh signalling genes may not be eliminated through cell competition, potentially causing abnormalities observed in diabetic pregnancies.

Furthermore, our findings suggest that acidosis (acidic pH stress) associated with maternal diabetes before or during pregnancy may contribute to increased susceptibility to conditions such as epilepsy, ASD, and later development of Alzheimer’s disease in offspring. Indeed, maternal diabetes elevates the risk of both epilepsy and ASD in humans^78,79^. Although evidence for Alzheimer’s disease in humans remains limited^80^, studies in mouse models showed that maternal high-fat diets can exacerbate Alzheimer’s-like symptoms in Tg offspring, including memory impairment and accumulation of phosphorylated tau^81,82^. Taken together, these observations suggest that in humans, pH stress caused by conditions such as diabetes impairs cell competition, allowing the accumulation of *de novo* mutant cells associated with epilepsy, ASD, and Alzheimer’s disease, thereby contributing to disease onset.

Developmental robustness refers to a system’s capacity to maintain precise embryonic development despite various perturbations. However, robustness paradoxically harbours a fragility to certain perturbations^83–85^. Nevertheless, little is known about the environmental factors that disrupt developmental robustness. Although ocean acidification reportedly causes abnormalities in embryonic development and organ formation in marine organisms, the underlying mechanisms are unknown^45–47^. Our study demonstrated that disruption of cell competition as a mechanistic link between environmental pH stress and developmental abnormalities, whereby acidic environments abolish cell competition and permit the accumulation of mutant cells with abnormal signalling, particularly affecting Wnt-dependent anterior-posterior patterning. Therefore, disruption of cell competition may be the mechanism underlying developmental abnormalities in marine organisms caused by ocean acidification.

Why are signalling-related genes disproportionately affected by *de novo* mutations? One explanation is their pivotal roles in axis formation, tissue patterning, and cell fate specification during early development, where even subtle perturbations can lead to pronounced developmental errors, rendering mutant cells more visible to the cell competition machinery. Another potential explanation for the enrichment of mutations in signalling-related gene sets is their high transcriptional activity and open chromatin state during early development. These features may render such loci more susceptible to DNA replication errors or DNA damage. Consistent with this idea, previous studies reported that highly transcribed regions with accessible chromatin tend to accumulate mutations due to impaired repair or increased exposure to genotoxic stress^86,87^. These observations suggest that signalling genes are not only functionally critical but also structurally prone to mutation during rapid embryogenesis. We also analysed copy number variations (CNVs); however, no clear differences were observed between control and cell death blocked larvae, likely reflecting the limited sensitivity of bulk-pooled WGS for low-frequency or mosaic CNVs. A single-cell WGS strategy will be crucial to map the full spectrum of mutations and chromosomal alterations subject to cell competition under physiological conditions. Moreover, because the analysis was performed using larvae rather than embryos, our dataset may be biased toward mutations that allow clonal expansion, whereas those impairing cell proliferation or survival could have been underrepresented. Nonetheless, our WGS results provide important evidence that endogenous cell competition plays a critical role in eliminating *de novo* mutant cells during development.

## Methods

### Maintenance of zebrafish

Zebrafish were raised and maintained under standard condition (14 h light/10 h dark cycle at 28.5). Wild-type strains (AB) were used along with the following transgenic lines: Tg(OTM:ELuc-CP)^12^. All experimental animal care was performed in accordance with the institutional and national guidelines and regulations. The study protocol was approved by the Institutional Animal Care and Use Committee of the University of Osaka (RIMD Permit# Biken-AP-R02-04).

### Preparation of plasmids

To prepare heat-shock promoter-driven plasmids, the *hsp70l* promoter was subcloned into the pTol2 vector (a gift from Dr. K. Kawakami). Subsequently, membrane-tagged (GAP43-fused) GFP (or mKO2) and T2A were subcloned into the pTol2-hsp70l promoter plasmids. These plasmids expressed only membrane-specific GFP (or mKO2) in response to heat shock. To generate plasmids expressing membrane GFP (or mKO2) with Wnt signalling genes, polymerase chain reaction (PCR)-amplified cDNAs encoding Wnt signalling regulators were subcloned downstream of T2A in pTol2-hsp70l:GFP-T2A (or mKO2-T2A) plasmids. The Wnt signalling activator was N-terminus-truncated mouse β-catenin (β-catCA), and the Wnt signalling inhibitor was human wild-type GSK-3β (a gift from Dr. A. Kikuchi).

For mRNA synthesis, cDNAs were PCR-amplified and cloned into the multi-cloning site of the pCS2p^+^ vector. Cloned cDNAs were as follows: human *bcl-2* (a gift from Dr. S. Korsmeyer, Addgene #8768)^88^; zebrafish DNA polymerase δ mutant (Polδ D406A), in which exonuclease domain Asp406 was substituted with Ala; zebrafish DNA polymerase ε mutant (Polε P285R), in which exonuclease domain Pro285 was substituted with Arg; Emerald luciferase^12^; H2B-fused GFP (or mCherry); GAP43-GFP (or mCherry); mCherry-SEpHluorin (a gift from Sergio Grinstein, Addgene #32001)^53^; secretory signal^89^ fused mCherry-SEpHluorin; and Ca^2+^ probe GCaMP7 (a gift from Koichi Kawakami)^90^.

Probes for *in situ* hybridization were *tbx6*^91^, *xirp2a*^92^, and *Eluc*^12^, which were subcloned into the pBluescript or pCRII TOPO or ppCS2p^+^ vector.

### mRNA synthesis and microinjection

Capped mRNA was synthesized using linearized plasmid DNA and an SP6 mMessage mMachine kit (AM1340; Thermo Fisher Scientific, Waltham, MA, USA). Synthesized mRNA was injected at the one-cell stage or the 8–16 cell stage of zebrafish embryos.

### Mosaic introduction of Wnt-abnormal cells

Hsp70 promoter-driven plasmids (5–17.5 pg) were injected into one-cell-stage embryos and maintained at 28.5°C until 4.3 hours post-fertilization (hpf) (dome stage). Subsequently, the embryos were exposed to heat shock. Briefly, the embryos were transferred to pre-warmed egg water at 37°C and kept at 37°C for 1 h. After heat shock, embryos were placed at 28.5°C, then fixed at 9 hpf for immunostaining or *in situ* hybridization. This method allowed the introduction of abnormal cells at the single-cell level, but not at the patchy-clone level.

### Mosaic introduction of polymerase mutant cells

Polδ D406A and Polε P285R mRNA with H2B-GFP and GAP43-GFP mRNA were injected into 8–16-cell-stage embryos (for mosaic expression) or one-cell-stage embryos (for ubiquitous expression), then maintained at 28.5°C and fixed at 9 hpf for immunostaining.

### sgRNA synthesis

We selected target sequences for each gene that did not overlap with other genomic sequences using the CRISPR design tool CHOPCHOP^93^. The oligonucleotides containing a T7 promoter sequence, target sequence, and the single-guide RNA (sgRNA) templates were PCR-amplified from sgRNA scaffold oligo using the oligonucleotides and primer sgRNA-Rv with KOD One (Toyobo, Osaka, Japan) and purified using the NucleoSpin Gel and PCR Clean-up Kit (MACHEREY–NAGEL, Düren, Germany). sgRNAs were synthesized using the CUGA7 gRNA Synthesis Kit (Nippon Gene, Tokyo, Japan).

### Mosaic or ubiquitous introduction of mutant cells using CRISPR/Cas9

Cas9 mRNA and sgRNAs were injected into a single blastomere of an 8- or 16-cell-stage embryo to introduce mosaic mutants. Cas9 mRNA and sgRNAs were injected into one-cell-stage embryos to generate ubiquitous mutants. Four sgRNAs were injected to disrupt the target gene efficiently^13^. The sgRNA target sequences were as follows: *appb*, 5′-gttgaagaagtatatccgtgcgg-3′, 5′-caacgtccagagtggcaagtggg-3′, 5′-aagcccaccgagtcatccgacgg-3′, 5′-cagagtgattacgatgacagtgg-3′; *psenen*, 5′-gggtggccgttttgacgacgtgg-3′, 5′-taaagaggccttcttaaaaccgg-3′, 5′-ctaaataatatctcctgcagagg-3′, 5′-gaggaatgcgaatccacctgagg-3′. As a control, four luciferase sgRNAs were injected; their sequences were as follows: 5′-gggcatttcgcagcctaccgtgg-3′, 5′-ggcatgcgagaatctcacgcagg-3′, and 5′-tcggggaagcggttgccaagagg-3′, and 5′-tttgtggacgaagtaccgaaagg-3′.

### Preparation of pH egg water

Zebrafish embryos were typically maintained in egg water (0.03% “SEALIFE” sea salt in water, MARINETECH, Tokyo, Japan). Normal pH egg water was prepared by adding HEPES (final concentration 10 mM; Dojindo, Kumamoto, Japan; #342-01375) and PIPES (final concentration 10 mM; Nacalai Tesque, Kyoto, Japan; #31212-25) and adjusting the pH to 7.0 with NaHCO_3_ (Nacalai Tesque; #31212-25). Acidic pH egg water was prepared by adding HCl (Nacalai Tesque; #18320-15), lactate (Sigma-Aldrich, St. Louis, MO, USA; #252476), or acetate (Nacalai Tesque; #00211-95) to the neutral pH egg water. Alkaline pH egg water was prepared by adding NaHCO_3_ to the neutral pH egg water.

### Antibodies

Primary antibodies were as follows: anti-γH2AX (#2577, 1/500 dilution; Cell Signalling Technology, Danvers, MA, USA); anti-GFP (#ab13970, 1/2000; Abcam, Cambridge, UK); anti-active caspase-3 (#9661, 1/500; Cell Signalling Technology); anti-mKO2 (#M168-3M, 1/500; Medical & Biological Laboratories, Tokyo, Japan); anti-mCherry (#ab125096, 1/500; Abcam); and anti-E-cadherin (#610181, 1/200; BD Biosciences, Franklin Lakes, NJ, USA). Secondary antibodies were as follows: AlexaFluor488-conjugated anti-mouse IgG (#A-11029, 1/500; Thermo Fisher); anti-rabbit IgG (#A-11034, 1/500; Thermo Fisher); anti-chicken IgY (#A78948, 1/2000; Thermo Fisher); AlexaFluor594-conjugated anti-mouse IgG (#A-11032, 1/500; Thermo Fisher); anti-rabbit IgG (#A-11037, 1/500; Thermo Fisher); and AlexaFluor647-conjugated anti-rabbit IgG (#4414, 1/500; Cell Signalling Technology).

### Chemical treatments

To induce Ca^2+^ influx, zebrafish embryos were treated with the Ca^2+^ ionophore ionomycin (110–330 nM; Cayman Chemical, Ann Arbor, MI, USA, #10004974).

### Whole-mount immunostaining

Embryos were fixed with 4% paraformaldehyde in phosphate-buffered saline (PBS) overnight at 4°C. Dechorionated embryos were washed four times with 0.5% Triton X-100 in PBS (PBST) and subsequently blocked with 10% fetal bovine serum, 4% Block Ace (Megmilk Snow Brand, Tokyo, Japan), and 1% dimethyl sulfoxide in 0.1% PBST for 1 h. The embryos were incubated with the primary antibodies overnight at 4°C, washed with PBST, and then incubated with AlexaFluor-conjugated secondary antibodies overnight at 4°C. Stained embryos were visualised with an M205FA fluorescence stereomicroscope (Leica, Wetzlar, Germany), a Thunder imager 3D (Leica), and an FV3000 confocal laser-scanning microscope (Olympus, Tokyo, Japan). Images were prepared and analysed using ImageJ software (NIH, Bethesda, MD, USA) or Aivia software (Leica).

### Whole-mount *in situ* hybridization

Whole-mount *in situ* hybridization was performed according to a standard protocol. Fluorescence *in situ* hybridization was performed as previously described^12^. Digoxigenin- or fluorescein isothiocyanate-labelled RNA antisense probes were prepared from plasmids containing *Eluc*, *tbx6*, and *xirp2a*. Images were captured using an M205A stereomicroscope (Leica).

### Time-lapse imaging of Ca^2+^ influx and data analysis

For time-lapse confocal live imaging, zebrafish embryos were manually dechorionated using forceps and mounted in 1% low-melting agarose in egg water on glass-bottom dishes. Live imaging was performed using an FV3000 confocal laser scanning microscope with excitation at 488 and 561 nm. Images were acquired every 10 s for 15 min to monitor Ca^2+^ dynamics using the GCaMP7 calcium indicator. Collected image data were analysed using ImageJ software. GCaMP7 fluorescence intensity (ΔF/F) was calculated by normalizing each time point to the mean fluorescence intensity at the initial (t = 0) and final (t = last) time points, and the results were plotted.

### Whole-genome sequencing analysis

*bcl-2* mRNA (200 pg) was injected into zebrafish embryos at the one-cell stage. Emerald luciferase (ELuc) mRNA served as a negative control. Four larvae injected with *bcl-2* mRNA at 32 hpf were analysed for the *bcl-2*-injected group (n = 3). The larvae were then snap-frozen in liquid nitrogen. Genomic DNA was extracted using a Genomic-tips 20/G kit (Qiagen, Hilden, Germany; #10223). Whole-genome sequencing (150 bp paired-end; 30× coverage) was performed by Relixa, Inc. (Tokyo, Japan). The extracted DNA was used to prepare paired-end libraries using the Truseq DNA PCR-free Library Prep Kit/Truseq DNA nano. Sequencing was conducted using a NovaSeq 6000 (Illumina, San Diego, CA, USA). The quality of raw paired-end sequence reads was assessed using FastQC (v0.11.7)^94^. Low-quality (<20) bases and adapter sequences were trimmed using Trimmomatic software (v0.38)^95^ with the following parameters: ILLUMINACLIP: path/to/adapter. fa:2:30:10 LEADING:20 TRAILING:20 SLIDINGWINDOW:4:15 MINLEN:36. The trimmed reads were aligned to the indexed reference genome (GRCz11; GCA_000002035.4) using BWA-MEM (v0.7.17-r1188)^96^. SAM files were converted into BAM files and sorted using Picard SortSam (v2.18.11). The number of duplicates was calculated using Picard Mark Duplicates. After removing duplicates, single-nucleotide variants and short insertions/deletions were called using Freebayes software (v1.3.4)^97^. The identified variants were annotated using SnpEff (v4.3t)^98^ to identify putative effects on protein translation and high-impact mutations. CNVs were analyzed using CNVkit (v0.9.8)^99^. Gene Ontology analysis of the mutated genes in apoptosis-inhibited larvae was performed using ShinyGo v0.77^100^. Genes that exhibited an increase in variant allele frequency (VAF) in two or three of the three analysed *bcl-2*-injected larvae (n = 3) are listed in Fig2D and 2E.

### Statistical analysis

Differences between groups were examined using a Mann-Whitney test, one-way analysis of variance, and Fisher’s exact test in GraphPad Prism10 software (GraphPad, Inc., San Diego, CA, USA).

## Data availability

The sequence data generated in this study were deposited in the NCBI Sequence Read Archive (https://www.ncbi.nlm.nih.gov/sra) under BioProject PRJNA1073592.

## Acknowledgments

We thank K. Kawakami, A. Kikuchi, W. El-Deiry, and S. Korsmeyer for providing plasmids, and Ishitani lab members for their helpful discussions, technical support, and maintenance of fish.

## Funding

The Takeda Science Foundation (T.I.)

SECOM Science and Technology Foundation (T.I.)

The Nippon Foundation - Osaka University Project for Infectious Disease Prevention (T.I.)

Grant-in-Aid for Transformative Research Areas (A) (21H05287) (T.I.)

Grant-in-Aid for Challenging Exploratory Research (23K18242) (T.I.)

Grant-in-Aid for Scientific Research on Innovative Areas (22H04845) (Y.A.)

Grant-in-Aid for Scientific Research on Innovative Areas (22H04845) (Y.A.)

Grant-in-Aid for Scientific Research(C) (25K09630) (Y.A.)

Astellas Foundation for Research on Metabolic Disorders (2022A1209) (Y.A.)

MSD Life Science Foundation, Public Interest Incorporated Foundation (Y.A.)

OU Master Plan Implementation Project promoted under Osaka University (T.I).

AMED-CREST (24gm2010001h0001) (T.I.).

AMED-IRUD (24ek0109760s4201) (T.I.)

A3 Foresight program (JPJSA3F20230001) (T.I.)

MEXT Promotion of Development of a Joint Usage/Research System Project: Coalition of Universities for Research Excellence (CURE) Program (JPMXP1323015484/JPMXP1323015486) (T.I.),

JST SPRING (H.U.).

## Author contributions

Conceptualization: Y.A. and T.I.

Methodology: Y.A., H.U., and T.I.

Investigation: Y.A., H.U., and T.I.

Visualization: Y.A., H.U., and T.I.

Funding acquisition: Y.A. and T.I.

Project administration: Y.A. and T.I.

Supervision: T.I.

Writing – original draft: Y.A. and T.I.

Writing – review & editing: Y.A. and T.I.

## Competing interests

Authors declare that they have no competing interests.

## Data and materials availability

All data needed to evaluate the conclusions in the paper are present in the paper and the Supplementary Materials. Raw sequence data are available through NCBI Sequence Read Archive (PRJNA1073592).

---

**Fig. S1.**
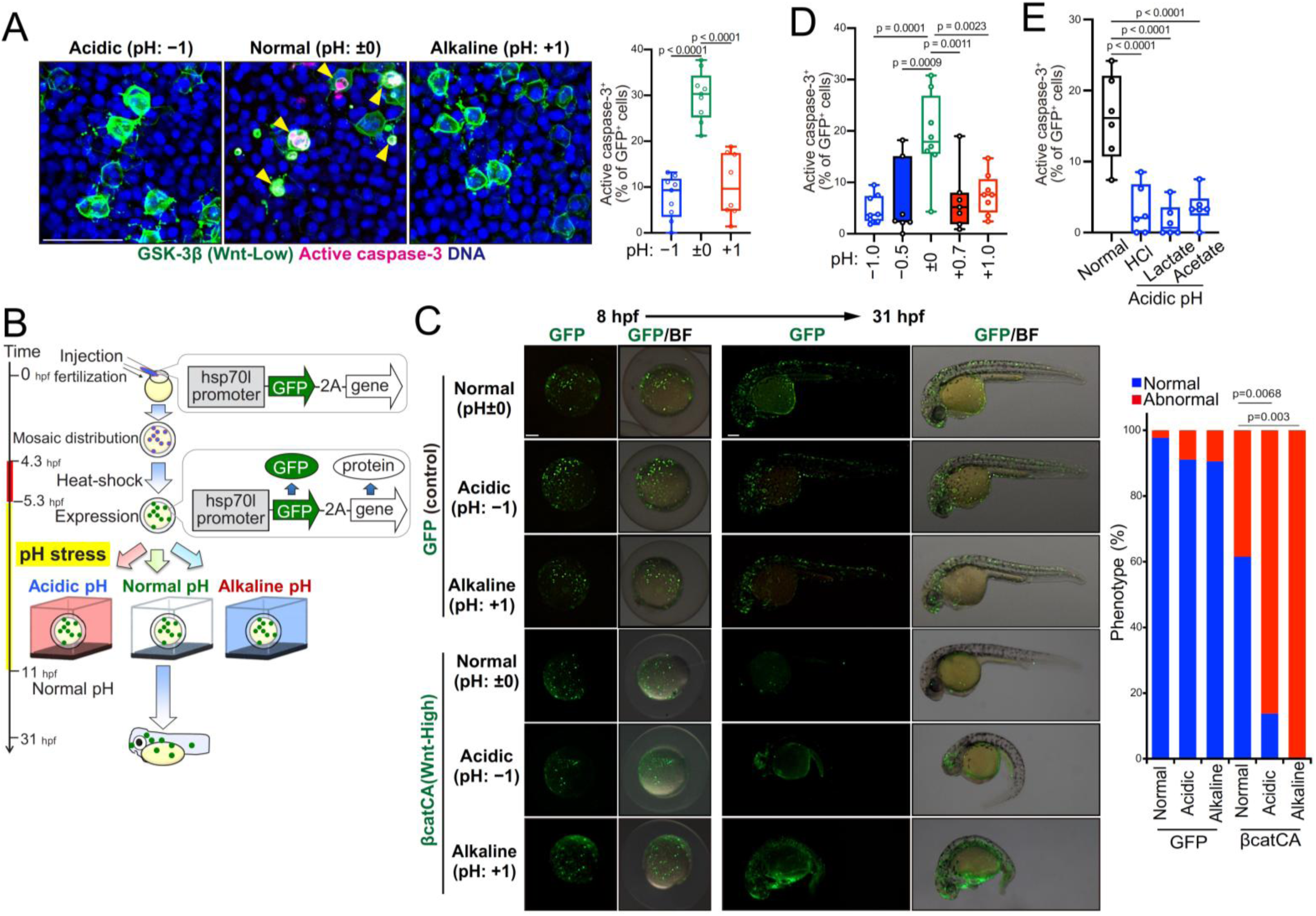
pH stress impairs artificially induced Wnt-driven cell competition, leading to abnormal development. (**A**) Artificially introduced GSK-3β-expressing (Wnt-Low) cells undergo apoptosis in a normal pH environment but not in acidic and alkaline pH environments. Confocal images show whole-mount immunostaining of active caspase-3 and DNA. Scale bars, 50 μm. Arrowheads indicate active caspase-3^+^ GSK-3β-expressing cells. Box plots show the percentage of GSK-3β-expressing cells (Wnt-Low) that are active caspase-3^+^, calculated by dividing the number of double-positive cells by the total number of GSK-3β-expressing cells; each dot represents one embryo. In total, approximately 400 GSK-3β-expressing cells were quantified per condition across embryos. (**B**) Schematic illustration of the experimental introduction of fluorescently labelled Wnt signal-abnormal cells into zebrafish embryos in a mosaic manner through heat shock induction. After heat shock (5.3 hour post-fertilization [hpf]), embryos were treated with the respective pH water, returned to normal pH at 11 hpf, and observed at 8 hpf and 31 hpf. (**C**) pH stress inhibits cell competition, leading to an accumulation of cells with abnormal Wnt signalling, resulting in developmental abnormalities. Images show embryos (8 hpf) and larvae (31 hpf) introduced with fluorescently labelled Wnt signalling aberrant cells (βcatCA-expressing cells) in a mosaic manner. Heat shock for mosaic induction was performed from 4.3 to 5.3 hpf. pH stress was applied from 5.3 to 11 hpf, before the return to normal water. The graph displays percentages of embryos with normal or abnormal morphology. (**D**) Inhibition of cell competition was also observed under milder acidic/alkaline environmental conditions. Box plots show the percentage of βcatCA-expressing cells that are active caspase-3^+^, calculated by dividing the number of double-positive cells by the total number of βcatCA-expressing cells; each dot represents one embryo. In total, approximately 400 βcatCA-expressing cells were quantified per condition across embryos. (**E**) In addition to hydrochloric acid, acidic pH environments prepared with lactic acid and acetic acid, which are abundant in tumour microenvironment, similarly suppressed Wnt-driven cell competition. Box plots show the percentage of βcatCA-expressing cells that are active caspase-3^+^, calculated by dividing the number of double-positive cells by the total number of βcatCA-expressing cells; each dot represents one embryo. In total, approximately 300-400 βcatCA-expressing cells were quantified per condition across embryos.

**Figure S2.**
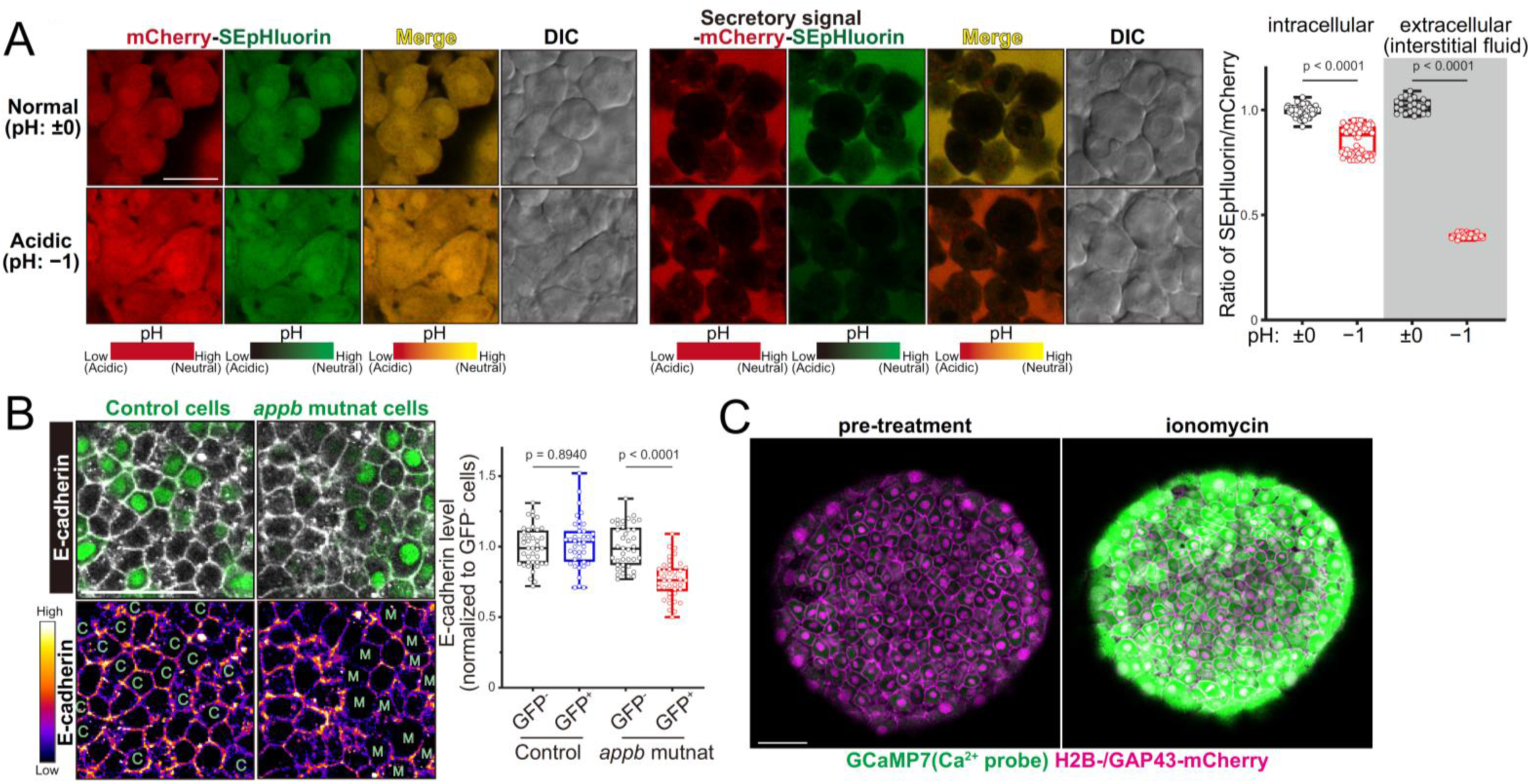
Supplementary evidence for pH stress-induced impairment of cell competition via dysregulated Ca²⁺ influx. (**A**) Extracellular pH shifts substantially under acidic conditions, whereas intracellular pH only changed slightly. Confocal images show zebrafish embryo expressing pH sensor (mCherry-SEpHluorin and secretory signal fused mCherry-SEpHluorin) under normal (pH: ±0) or acidic (pH: -1.0) condition in egg water. Scale bar, 10 μm. Box plots indicate the relative SEpHluorin/mCherry ratio in the intracellular or extracellular (interstitial fluid), normalized to the normal pH condition. (**B**) E-cadherin levels are downregulated in amyloid β precursor protein b (*appb*) mutant cells. Confocal images show immunostaining of E-cadherin in embryos mosaically introduced with control sgRNAs or sgRNA targeting *appb* together with Cas9. C, control cell; M, mutant cell. Scale bar, 50 μm. (**C**) Ionomycin induces forced Ca^2+^ influx. Confocal images of embryos expressing the Ca^2+^ probe GCaMP7 and nuclear/membrane-localized mCherry (H2B-/GAP43-mCherry), shown before (pre-treatment) and after (post-treatment) ionomycin exposure. Scale bar, 50 μm.

**Movie S1 pH stress induces chaotic Ca^2+^ influx**

Zebrafish embryo injected with mRNA encoding the Ca^2+^ probe GCaMP7 and nuclear/membrane-localized mCherry (H2B-/GAP43-mCherry), exposed to egg water at normal, acidic, or alkaline pH. Images were acquired at 10-s intervals for 15 min. Scale bar, 50 μm.

**Movie S2 Validation of ionomycin-induced elevation of intracellular Ca^2+^ concentration**

Zebrafish embryo injected with mRNA encoding Ca^2+^ probe GCaMP7 and nuclear/membrane-localized mCherry (H2B/GAP43-mCherry), treated with ionomycin (330 nM) following a pre-treatment period. Images were acquired at 10-s intervals for 30 min. Scale bar, 50 μm.

## Notes

### Competing Interest Statement

The authors have declared no competing interest.

